# Target membrane cholesterol modulates single influenza virus membrane fusion efficiency but not rate

**DOI:** 10.1101/2019.12.19.882738

**Authors:** K. N. Liu, S. G. Boxer

**Affiliations:** Stanford University

## Abstract

Host lipid composition influences many stages of the influenza A virus (IAV) entry process, including: initial binding of IAV to sialylated glycans, fusion between the viral envelope and the host membrane, and the formation of a fusion pore through which the viral genome is transferred into a target cell. In particular, target membrane cholesterol has been shown to preferentially associate with virus receptors and alter physical properties of the membrane like fluidity and curvature. These properties affect both IAV binding and fusion, which makes it difficult to isolate the role of cholesterol in IAV fusion from receptor binding effects. Here, we develop a new fusion assay that uses synthetic DNA-lipid conjugates as surrogate viral receptors to tether virions to target vesicles. To avoid the possibly perturbative effect of adding a self-quenched concentration of dye-labeled lipids to the viral membrane, we tether virions to lipid-labeled target vesicles, and use fluorescence microscopy to detect individual, pH-triggered IAV membrane fusion events. Through this approach, we find that cholesterol in the target membrane enhances the efficiency of single-particle IAV lipid mixing, while the rate of lipid mixing is independent of cholesterol composition. We also find that the single-particle kinetics of influenza lipid mixing to target membranes with different cholesterol compositions is independent of receptor binding, suggesting that cholesterol-mediated spatial clustering of viral receptors within the target membrane does not significantly affect IAV hemifusion. These results are consistent with the hypothesis that target membrane cholesterol increases lipid mixing efficiency by altering host membrane curvature.

**Statement of Significance:** Influenza A virus is responsible for millions of cases of flu each year. In order to replicate, influenza must enter a host cell through virus membrane fusion, and cholesterol in the target membrane is vital to the dynamics of this process. We report a receptor-free, single virus fusion assay that requires no fluorescent labeling of virus particles. We use this assay to show that cholesterol increases the fraction of fusion events in a manner that is correlated with the spontaneous curvature of the target membrane but is independent of receptor binding. This assay represents a promising strategy for studying viral fusion processes of other enveloped viruses.

## Introduction

In order for influenza A virus (IAV) to replicate, hemagglutinin (HA), a viral envelope protein, must first bind to negatively-charged sialylated glycolipids or glycoproteins on the host plasma membrane. Once IAV is bound and endocytosed, viral content entry is facilitated by membrane fusion, where endosomal low pH triggers a conformational rearrangement of HA, and the membranes of IAV and the target cell mix to form a hemifusion intermediate (1, 2). Lipid mixing is followed by the formation of a fusion pore through which the viral genome is transferred into the target cell (3, 4).

Host membrane composition influences each stage of the IAV entry process, though it is not entirely clear how it does so on a molecular scale (5, 6). Target membrane cholesterol has been shown to cluster with sialic acid-presenting glycolipids, which increases IAV binding avidity (7). It has also been demonstrated in bulk studies that cholesterol in target vesicles speeds the rates of IAV hemifusion and fusion pore formation (8), which may be facilitated by the negative spontaneous curvature of cholesterol (9). For other enveloped viruses like human immunodeficiency virus (HIV), fusion peptide insertion is facilitated by cholesterol in the target membrane, which has led to speculation that cholesterol may play some role in facilitating HA engagement for IAV (10–12). Given that cholesterol may stabilize each fusion intermediate, bulk measurements cannot deconvolve the mechanistic details of cholesterol’s influence on IAV fusion.

Single-particle virus lipid mixing techniques have enabled researchers to separate the dynamics of virion binding from subsequent steps in IAV membrane fusion (13). These approaches typically utilize fluorescence microscopy and dequenching to monitor receptor-bound individual virions fusing to artificial membrane bilayers (14–16). Models and simulations based on experimental kinetics have shown that lipid mixing is the rate limiting step of IAV membrane fusion (17, 18). It is postulated that a minimum of 3 neighboring HA trimers must engage in the target membrane in order to yield a productive hemifusion event (17, 19). This estimate suggests that clustering of HAs, as a result of receptor binding and possibly mediated or affected by membrane composition, could enhance the rate of IAV fusion.

It has been difficult to isolate the role of receptor binding in IAV fusion because HA is responsible for both IAV binding and fusing to the host membrane. To disentangle receptor binding and fusion, our lab previously developed a single-particle assay that uses synthetic DNA-lipid surrogate receptors and lipid dequenching assays to observe lipid mixing between single virions and surface-tethered vesicles (20, 21) (Fig. S1 in the Supporting Material). Application of this assay demonstrated that kinetics of IAV lipid mixing to target membranes containing 10 mol% cholesterol are the same when virus and vesicle are tethered through DNA-lipid surrogates or GD1a sialic acid receptors.

Despite the fact that single-particle approaches have yielded an immense amount of kinetic information about IAV fusion, a limitation of currently existing strategies is that they all involve introducing a self-quenched concentration of fluorescently tagged lipids to the viral envelope. The labeling process exposes virions to organic solvents like ethanol and DMSO, which can potentially impact virus infectivity. At self-quenched concentrations, unnatural fluorescent lipids comprise roughly 3-5 mol% of the IAV envelope, and this can affect the observed lipid mixing kinetics of IAV fusion (22).

In this study, we introduce a new assay architecture to observe single-particle lipid mixing events, illustrated in Fig. 1 (compare to Fig. S1), that limits the fluorescent label to the target vesicle, thereby avoiding the process of adding fluorescent dye to viruses. As in our previous assay, this approach utilizes sequence-specific DNA-lipid conjugates to tether IAV to target membranes in the absence of the native sialic acid viral receptor. We apply this strategy to study the impact of target membrane cholesterol on IAV fusion kinetics at the single-particle level. By varying the amount of cholesterol in target membranes from 10-40 mol%, we show that while cholesterol enhances the efficiency, or fraction of productive IAV lipid mixing events, the rate of fusion is independent of target membrane composition, in both the presence and absence of the native viral receptor GD1a. This suggests that cholesterol-mediated spatial clustering of viral receptors in the target membrane does not have a significant impact on the rate limiting step of IAV lipid mixing. Additionally, we relate the cholesterol dependence of IAV lipid mixing efficiency to the spontaneous curvature of the target membrane.

**Figure 1.**
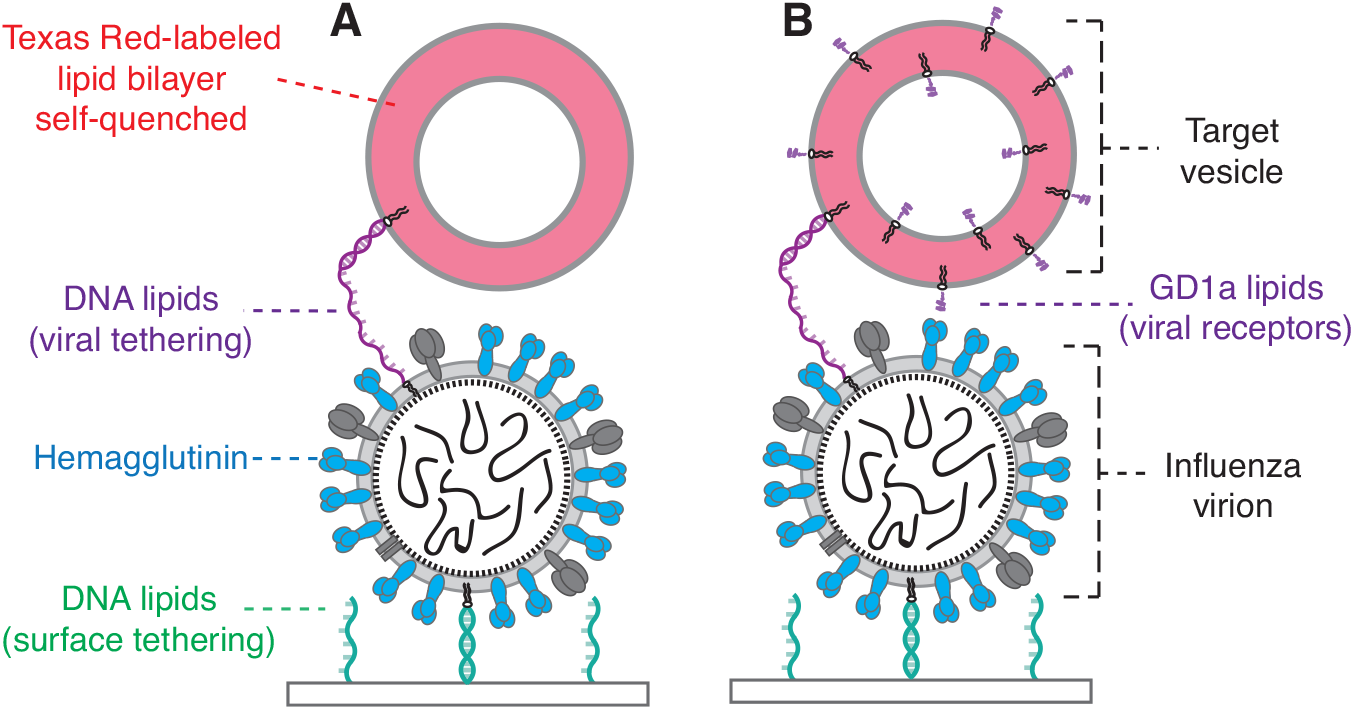
Schematic of influenza A virions bound to Texas Red lipid-labeled target vesicles. Vesicles (red) are bound to virions by (A) hybridization of complementary sequences of membrane-anchored DNA (purple) or (B) both membrane-anchored DNA and GD1a glycolipid receptors (black). GD1a binds to viral hemagglutinin (blue). In both architectures, the substrate displays single-stranded DNA (green). Virions are surface-tethered by DNA hybridization with a membrane-anchored antisense DNA sequence (green) that is orthogonal to the sequence used for viral tethering (purple). Details of surface functionalization not shown (see Supporting Materials); schematic not drawn to scale.

## Materials and Methods

### Materials

Palmitoyl oleoyl phosphatidylcholine (POPC), dioleoyl phosphatidylethanolamine (DOPE), and cholesterol were purchased from Avanti Polar Lipids (Alabaster, AL). Texas Red-1,2-dihexadecanoyl-*sn*-glycero-3-phosphoethanolamine (TR-DHPE) and Oregon Green-1,2-dihexadecanoyl-*sn*-glycero-3-phosphoethanolamine (OG-DHPE), fatty acid depleted bovine serum albumin (BSA), and NeutrAvidin were purchased from Thermo Fisher Scientific (Waltham, MA). Disialoganglioside GD1a from bovine brain (Cer-Glc-Gal(NeuAc)-GalNAc-Gal-NeuAc) was purchased from Sigma-Aldrich (St. Louis, MO). Chloroform, methanol, and buffer salts were obtained from Fisher Scientific (Pittsburgh, PA) and Sigma-Aldrich. Polydimethylsiloxane (PDMS) was obtained from Ellsworth Adhesives (Hayward, CA). Poly(L-lysine)-graft-poly(ethylene glycol) (PLL-g-PEG) and Poly(L-lysine)-graft-poly(ethylene glycol) biotin (PLL-g-PEG biotin) were purchased from SuSoS AG (Dübendorf, Switzerland).

### Buffers

The following buffers were used. Vesicle buffer: 10 mM NaH_2_PO_4_, 90 mM sodium citrate, 150 mM NaCl, pH 7.4. Fusion buffer: 10 mM NaH_2_PO_4_, 90 mM sodium citrate, 150 mM NaCl, pH 5.1. HB buffer: 20 mM HEPES, 150 mM NaCl, pH 7.2.

### Microscopy

Epifluorescence micrographs were acquired with a Nikon Ti-U microscope using a 100X oil immersion objective, NA = 1.49 (Nikon Instruments, Melville, NY), a Spectra-X LED Light Engine (Lumencor, Beaverton, OR) for illumination and an Andor iXon 897 EMCCD camera (Andor Technologies, Belfast, UK) with 16-bit image settings. Images were captured with Metamorph software (Molecular Devices, Sunnyvale, CA). See Supporting Material for additional microscopy information.

### DNA-lipid and biotin-DNA preparation

DNA-lipids (see Table S1 for sequences) used to surface-tether viruses and tether target vesicles to viruses were synthesized as previously described (23). Biotin-DNA was synthesized by the PAN facility at Stanford University and diluted to the desired concentration in DI water. All DNA oligos were stored at −20°C.

### Influenza virus preparation

Influenza A virus (strain X-31, A/Aichi/68, H3N2) was purchased from Charles River Laboratories (Wlimington, MA). Virus was pelleted in HB buffer by centrifugation at 21130 rcf for 50 min and resuspended in 100 μL of fresh HB buffer. DNA-lipids were incorporated into the IAV envelope by incubating virus sample at 4°C on ice overnight according to a previously described method (20). IAV is a BSL-2 agent and was handled following an approved biosafety protocol at Stanford University.

### Surface and architecture preparation

The single virus lipid mixing architecture was prepared as described in Fig. 1. In a microfluidic flow cell (see Supporting Material for details), glass slides were functionalized by incubating a mixture of 5 μL of a 19:1 mixture of PLL-g-PEG (1 g/L) and PLL-g-PEG biotin (1 g/L) in HB buffer for at least 30 min. Flow cells were rinsed with DI water and vesicle buffer, and stored overnight at 4°C. The following day, 5 μL of neutrAvidin (1 g/L) was incubated for 15 min. After rinsing away excess neutrAvidin with vesicle buffer, 2 μL of biotin-DNA (178 μM, sequence A – see Table S1 for sequences) was incubated for 20 min. The flow cell was thoroughly rinsed with vesicle buffer to remove excess biotin-DNA. Next, 5 μL of IAV in HB buffer (roughly 5.4 nM) displaying sequences A’ (antisense to A) and B were introduced to the flow cell and tethered to the substrate. After rinsing the flow cell with vesicle buffer, the surface was further passivated by incubating 10 μL of bovine serum albumin (BSA, 1 g/L) for at least 10 min to prevent non-specific binding of added target vesicles. Finally, 2-3 μL of ~100 nm diameter vesicles displaying DNA sequence B’ and/or containing GD1a were introduced (2.8 μM nominal total lipid concentration) and allowed to bind for 5-10 min to control surface density and ensure spatial separation between particles. Excess unbound vesicles were removed by rinsing with vesicle buffer.

### Lipid-mixing assay

Fluorescence microscopy was used to collect a video micrograph image stream of 1000 frames at a rate of 3.47 frames/s. After the image stream was started, the pH of flow cell was rapidly exchanged from 7.4 to 5.1 using fusion buffer. In a separate experiment, tethered vesicles that contained a pH indicator (2 mol% OG-DHPE) were used to calibrate the exchange time of fusion buffer (2-3 s). The time between lowering of pH to dequenching was extracted using custom MATLAB (MathWorks, Inc.) scripts, as described previously (20, 21). In 0.5% of traces or fewer, more than one dequenching event is observed in the same region, reflecting that more than one vesicle was bound to the same virion, or that there was insufficient spatial separation between particles. The wait times from fluorescence traces with more than one dequenching event are excluded from CDFs.

## Results and Discussion

### Single virus lipid mixing assay

We developed an assay that monitors single lipid mixing events and eliminates the need to fluorescently label virions (assay schematized in Fig. 1). In this strategy, synthetic DNA-lipids are used for two purposes: 1) to surface-immobilize influenza virions and 2) to tether target vesicles to virions in the presence and absence of sialic acid receptors.

First, DNA-lipids with two orthogonal sequences (Fig. 1, green and purple) are incorporated into the envelope of influenza A virions. This is accomplished by incubating virions with an aqueous suspension of DNA-lipids, that is, no organic solvents are used. For each sequence, the median number of DNA-lipids incorporated each virion is 7 (Fig. S2). As previously observed, when DNA-lipids are incubated with virions or lipid vesicles, DNA-lipid incorporation into the IAV envelope does not follow Poisson statistics (20, 24). The median number of DNA-lipids per virion is 200- to 350-fold fewer than the number of fluorescently tagged lipids added to the viral envelope in order to achieve a self-quenched concentration in alternate architectures. Virions are added to a microfluidic flow cell containing a carefully passivated substrate. The substrate displays a complementary DNA sequence to one of the DNA-lipid sequences on the virion (green), leading to surface immobilization of the virions through DNA hybridization.

After the flow cell is rinsed extensively, bovine serum albumin (BSA) is added to further passivate the substrate and prevent non-specific binding of incoming target vesicles. Effective surface passivation requires optimization to ensure that the target vesicles are specifically bound to virions. Target vesicles containing DNA-lipids are introduced to the flow cell at a dilute concentration and become tethered to virions through hybridization of complementary DNA sequences (purple). Vesicles are bound sparsely for sufficient spatial resolution to monitor single events. These target vesicles are lipid-labeled with a self-quenched concentration of TR-DHPE (7 mol%). They are ~100 nm in diameter to ensure that mixing and dilution with the viral envelope membrane, which is also ~100 nm diameter (albeit quite heterogenous (25)) will result in dequenching. In the absence of IAV or sequence-specific DNA-tethers, very few vesicles bind to the passivated substrate. Unbound vesicles are thoroughly rinsed and removed from the flow cell.

To study lipid mixing kinetics in the presence of sialic acid receptors, target vesicles were also prepared with sialic acid-containing glycolipids (ganglioside GD1a) in addition to DNA-lipids. Due to the transient and reversible interaction of IAV with GD1a, DNA-lipids are required during preparation of target vesicles to ensure that vesicles remain bound.

Once virions and target vesicles are bound, the pH of the flow cell is lowered from pH 7.4 to 5.1 through rapid buffer exchange. A video micrograph monitoring the Texas-Red signal of vesicles is collected for 1000 frames at 288 ms/frame. The dequenching of Texas-Red DHPE represents lipid transfer from the target vesicle to the IAV envelope (Fig. 2A and B). Fluorescence traces of individual events are analyzed to extract the time interval from pH drop to lipid mixing (Fig. 2C). Individual wait times are compiled in a cumulative distribution function (CDF, Fig. 3A and B). CDFs were used rather than histograms to present wait times because they do not require data to be selectively binned. In the absence of IAV, there are no vesicles that undergo fluorescence dequenching when the pH of the system is lowered (Fig. S3), which indicates that all vesicle dequenching events observed can be attributed to viral lipid mixing. The kinetics of lipid mixing observed with this architecture are indistinguishable from our previously published single virus measurements that monitored dequenching of lipid-labeled IAV (20) (Fig. S4).

**Figure 2.**
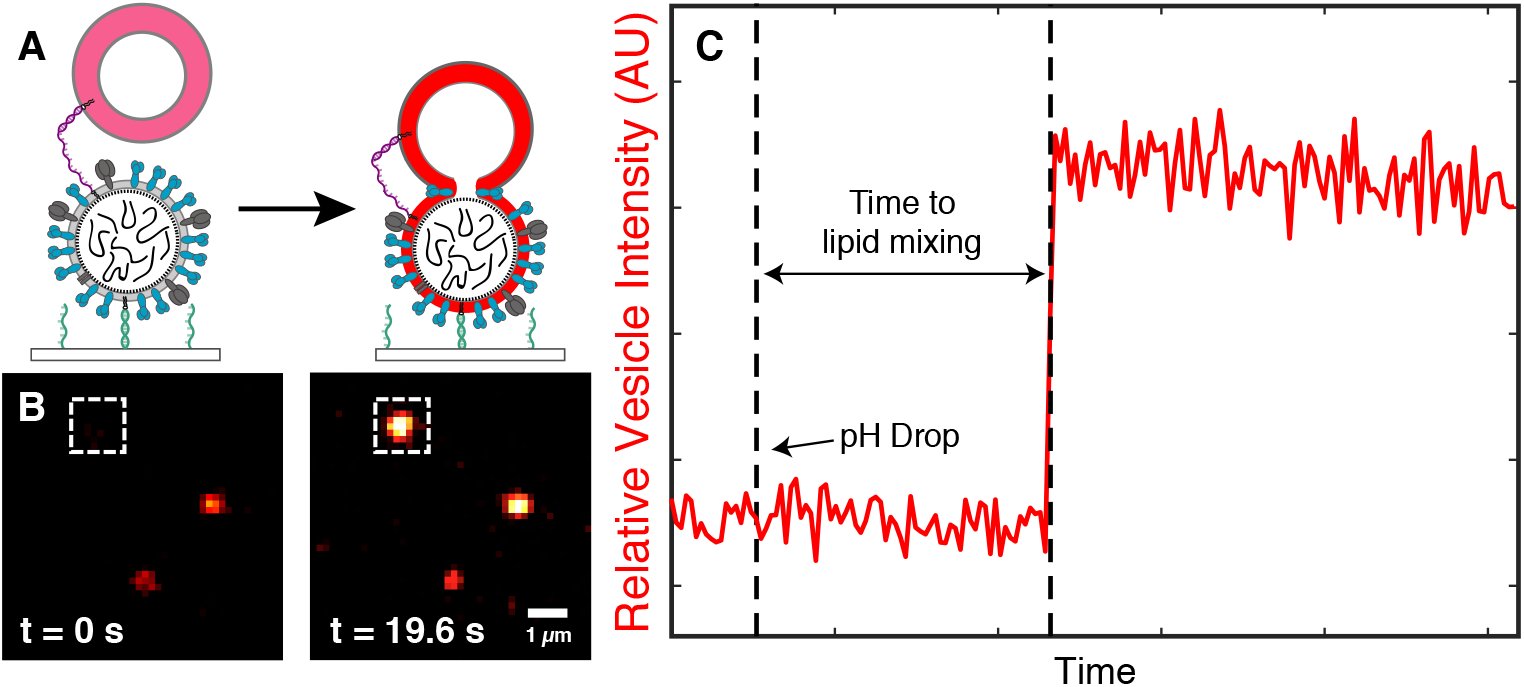
Lowering the pH triggers virion-target vesicle hemifusion detected by fluorescence dequenching. (A) Schematic diagram illustrating the single virus lipid mixing experiment. A Texas-Red labeled vesicle (self-quenched concentration) is tethered *via* DNA-lipids to a surface-tethered influenza virion (see Fig. 1A). When the pH is lowered, hemifusion occurs and the lipids in the target vesicle are diluted upon mixing with the viral membrane. Schematic is not drawn to scale. (B) Example microscope images of Texas-Red labeled vesicles that are bound to surface-tethered influenza virions. At pH 7.4 (left), the self-quenched target vesicle is dim but detectable. After the pH is lowered to 5.1 (right), particles display dequenching due to lipid mixing. (C) Example time trace (red) of the fluorescence intensity of a Texas-Red vesicle (shown in white box in B) that undergoes dequenching. The wait time is defined as the time from the pH drop to the lipid mixing event.

**Figure 3.**
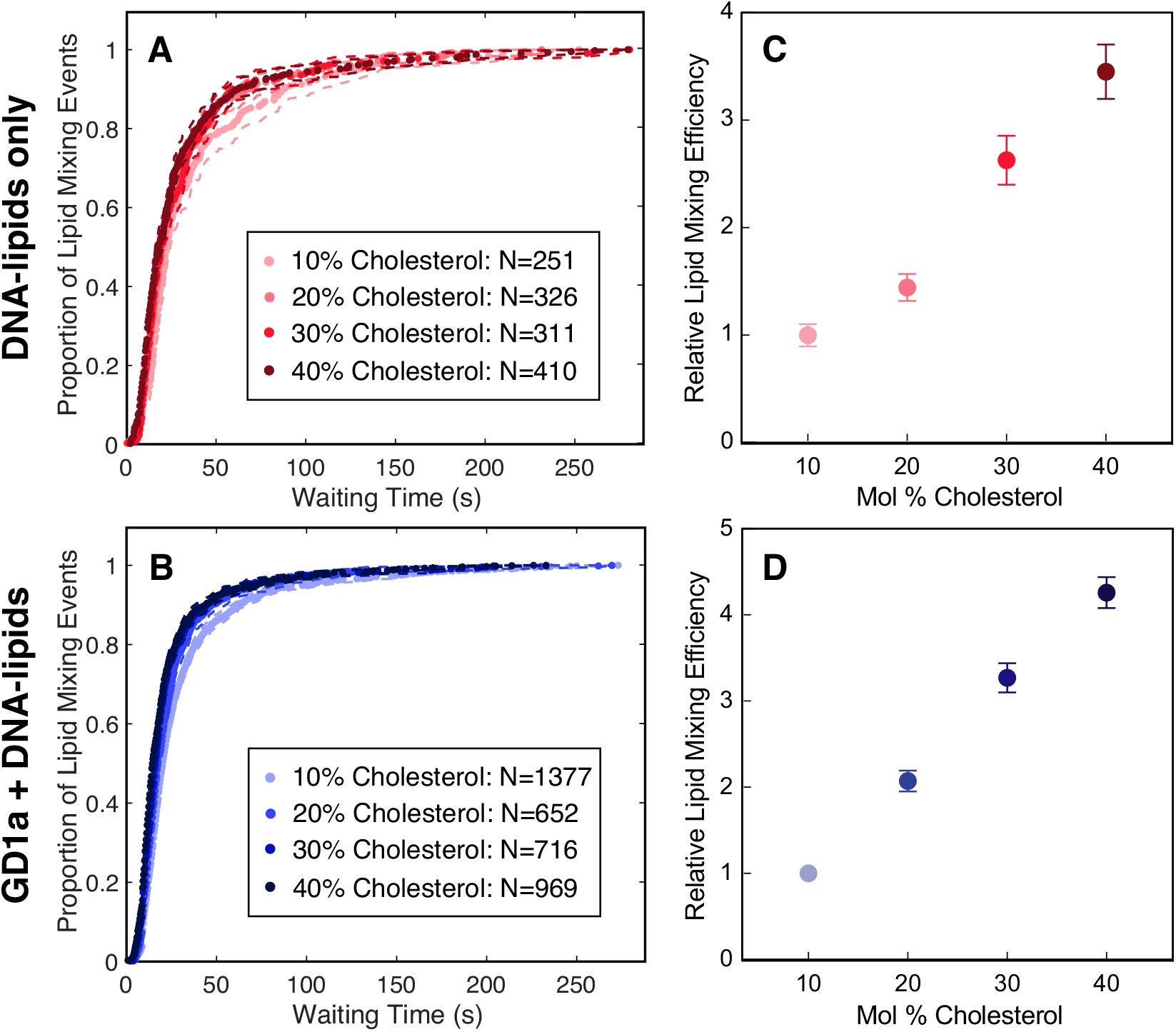
Cholesterol enhances the efficiency of single-particle IAV lipid mixing, while the rate of lipid mixing is independent of target membrane cholesterol composition. (A) Wait times for individual lipid mixing events are plotted as cumulative distribution functions (CDFs). From 10-40 mol% cholesterol, the rate of IAV lipid mixing is the same within bootstrap resampling error (95% confidence intervals, dashed lines; see Fig. S5 for individual CDFs). (B) In the presence of native viral receptor GD1a, the rate of lipid mixing is also unaffected by increasing the ratio of cholesterol in the target membrane within bootstrap resampling error (95% confidence intervals, dashed lines). (C) The relative efficiency or fraction of target vesicles that undergo dequenching increases 4-fold as the mol% of cholesterol in DNA-tethered target vesicles increases. For C and D, the lipid mixing efficiency to membranes containing 10 mol% cholesterol is normalized to 1, and points represent the average efficiency value ± bootstrap resampling error. (D) When target membranes contain GD1a, the relative efficiency of lipid mixing also increases 4-fold. Kinetic data for each composition were compiled from at least two independent viral preparations.

### Cholesterol enhances the efficiency of IAV lipid mixing, but does not alter the rate

To gain mechanistic insight into the role of cholesterol in IAV hemifusion, we performed single virus lipid mixing experiments on virions that were DNA-tethered to target membranes that contained varying mole fractions of cholesterol. The wait times to lipid mixing for each composition were compiled into separate CDFs (Fig. S5) and compared. Surprisingly, from 10-40 mol% cholesterol, the rates of IAV lipid mixing are the same within experimental error (Figure 3A). However, the relative efficiency of lipid mixing, or the fraction of vesicles within a field of view (FOV) that undergo dequenching, increases 4-fold as the target membrane cholesterol composition was increased from 10-40 mol% (Fig. 3C).

To understand whether cholesterol-mediated GD1a spatial ordering impacts the kinetics of lipid mixing, we introduced GD1a (2 mol%) to target vesicles in addition to DNA-lipids, then varied the mole fraction of cholesterol in the target membrane from 10-40 mol%. Similar to IAV lipid mixing with DNA-tethered target membranes, when GD1a is also present in target vesicles, the rates of IAV lipid mixing are independent of mole fraction of cholesterol in the target membrane (Fig. 3B). Increasing the amount of cholesterol from 10-40 mol% also enhances the relative efficiency of IAV lipid mixing by 4-fold (Fig. 3D).

Although target vesicles were bound by both GD1a and DNA-lipids, the quantity of GD1a glycolipids is roughly 40-fold greater than the median number of DNA-lipids per vesicle. Due to the large excess of GD1a, we hypothesize that the lipid mixing kinetics observed are primarily influenced by the GD1a-hemagglutinin interactions between IAV and target vesicle. Additionally, the rates found through the vesicle-labeled architecture in this study are indistinguishable from previous measurements of IAV-labeled, GD1a-mediated lipid mixing (20).

The single virus lipid mixing data from our study show that while cholesterol increases the fraction of productive hemifusion events, the rate of IAV lipid mixing is independent of cholesterol composition. These observations are consistent with the increase in extent of IAV fusion found in bulk ensemble measurements when target membrane cholesterol is increased (8, 26). In a different single-particle study, it was found that cholesterol increases both the rate and efficiency of lipid mixing of lipid-labeled influenza virions bound to target membranes by GD1a (27). We hypothesize that our results may differ from this observation due to differences in virion labeling, for example, the perturbative effect of adding large quantities of octadecyl rhodamine B (R18) to the viral envelope (22).

The close similarity of kinetics of IAV lipid mixing to receptor (GD1a)-containing and receptor-free target membranes supports our previous finding that the absence of GD1a does not impact the spatial organization of viral HA in a manner that alters the kinetics of hemifusion (20). This study further demonstrates that lipid mixing kinetics are independent of receptor binding across a range of target membrane lipid compositions. We do not directly measure HA engagement with the target membrane, so at this point we are unable make any claims about the impact of cholesterol on lipid mixing intermediates. Although cholesterol has been shown to cluster with target receptors (7, 28), the concentration of cholesterol in target membranes does not significantly alter the rate-limiting step of IAV fusion.

### Cholesterol may stabilize hemifusion to increase IAV lipid mixing efficiency

We sought to understand how cholesterol acts to modulate the physical properties of the target membrane to yield a greater number of productive IAV hemifusion events. In addition to the fact that cholesterol increases bilayer rigidity and leads to greater orientational order (29), cholesterol generates negative curvature in lipid bilayers, which can promote the formation of highly curved membrane structures. It is well established that the hemifusion stalk formed during vesicle (30) and IAV membrane fusion is stabilized by negative spontaneous curvature (31). Cryo-electron tomography images have shown that increasing cholesterol in the target membrane, or modulating spontaneous curvature through lipid composition, can induce different hemifusion structures and pathways to IAV fusion (32).

To understand the relationship between lipid mixing efficiency and curvature, we estimated the monolayer spontaneous curvature (SC) for each mixture tested in this study using published values for each individual membrane component (33) (Supporting Material, Table S2).Several assumptions were needed to estimate the lipid mixture SC. The calculations assume no asymmetry between inner and outer leaflet of target vesicles, although it is important to note that any asymmetry in lipid composition between membrane leaflets can impact the effective membrane SC. The contributions of TR-DHPE and ganglioside GD1a were also omitted. There is no agreed upon experimental value for the individual contribution of GD1a to membrane SC, although gangliosides like GM1 have been suggested to induce positive curvature (34), and GD1a has been shown to induce high curvature clusters in phosphatidylethanolamine (POPE) membranes (35). With these assumptions, for mixtures containing 10, 20, 30, and 40 mol% cholesterol, the estimated SC values were −0.142, −0.190, −0.237, and −0.284 nm^−1^, respectively.

As shown in Fig. 4, the lipid mixing efficiency values increase as target membrane SC becomes more negative. The relative efficiencies of lipid mixing increase slightly for virions that are both DNA-tethered and bound by GD1a, perhaps because GD1a may additionally induce curvature in these specific lipid mixtures that favorably stabilizes the hemifusion stalk. Another explanation is that in the absence of GD1a, DNA-tethering could potentially bind vesicles to a greater proportion of fusion-incompetent virions that are unable to bind to GD1a, therefore decreasing the relative efficiency. These two hypotheses are difficult to test and deconvolve as there is as of yet no way to directly characterize each individual virion (and target vesicle) whose kinetics are observed.

**Figure 4.**
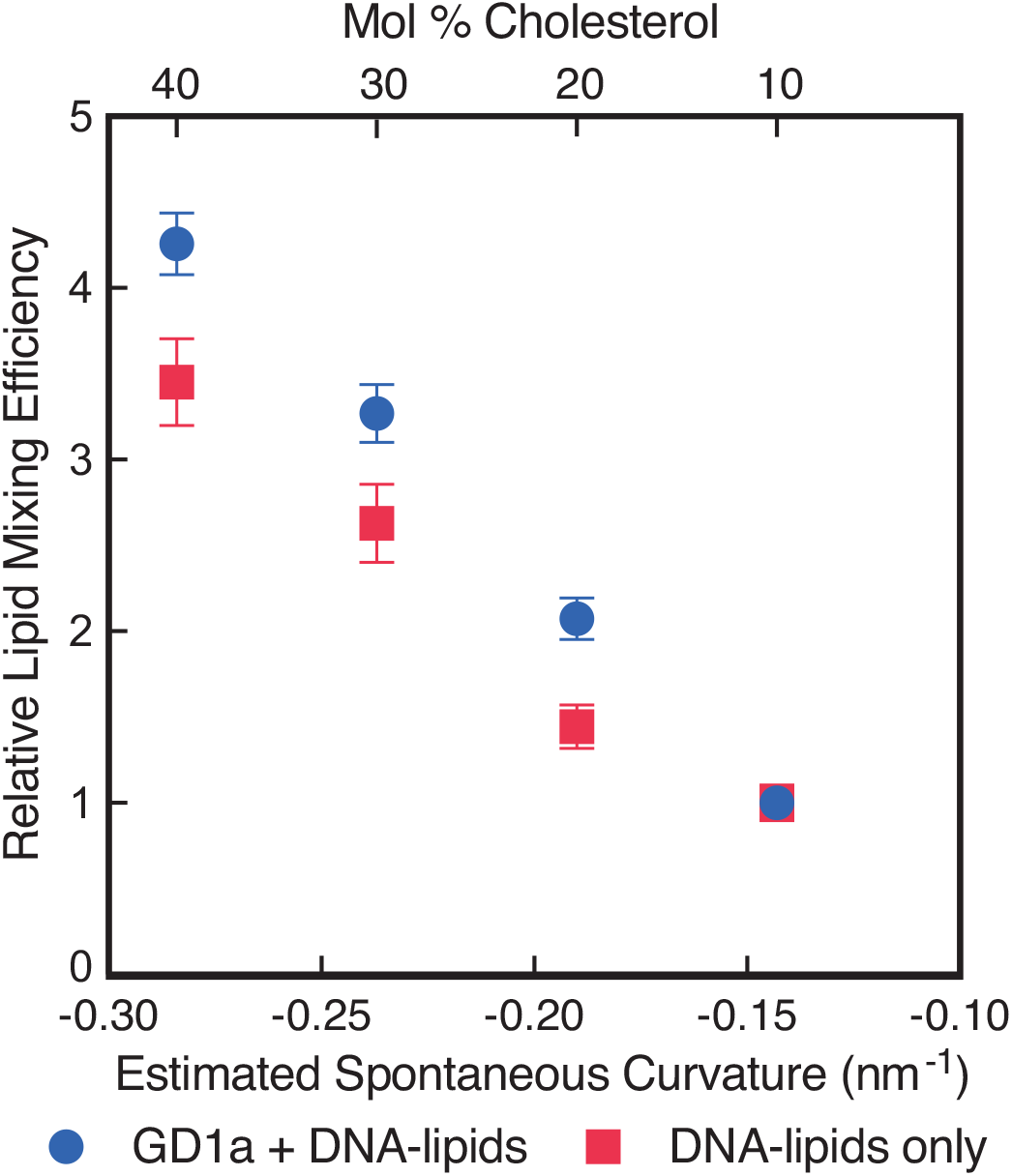
IAV lipid mixing efficiency increases when estimated negative spontaneous curvature becomes more negative. This trend is observed in both the presence (blue) and absence (red) of viral ganglioside receptor GD1a. Points represent the average efficiency value ± bootstrap resampling error.

Our analysis focuses on the curvature of the hemifusion intermediate formed between IAV and target membrane. Although the overall curvature of vesicles in this model system differs from the curvature of the endosomal membrane that IAV typically fuses to in a cell, the curvature of the hemifusion stalk in both environments is likely to be the same.

The correlation between target membrane SC and IAV lipid mixing efficiencies suggests that the dominant effect of cholesterol in IAV lipid mixing is in facilitating the creation of the hemifusion stalk intermediate. The stabilization of this fusion intermediate enables a greater number of successful lipid mixing events to occur, while imparting no significant perturbation on the rate-limiting step of IAV fusion.

## Conclusions

We have presented a strategy to observe single virus lipid mixing events that eliminates the need to add high concentrations of fluorescently-tagged lipids to the viral envelope. Using this approach, we find that the rate of IAV lipid mixing is insensitive to the mole fraction of cholesterol in the target membrane, while the efficiency is enhanced in a manner that is correlated with the spontaneous curvature of the target membrane. We also show that nanoscale clustering of GD1a and cholesterol in the target membrane does not have a significant impact on either the rate or relative efficiency of IAV lipid mixing.

While this study only reports lipid mixing of IAV, the general strategy of restricting fluorescent labels to the target vesicle is promising for observing other viral processes, in particular, content mixing, and this will be reported separately. This general architecture could also be adapted to observe target membrane fusion kinetics of other enveloped viruses, especially for viral samples that require harsh conditions to fluorescently label or where the receptor is either unknown or difficult to work with.

## Author Contributions

K.N.L. designed experiments, performed experiments, analyzed data, and co-wrote the article. S.G.B. designed experiments and co-wrote the article.

## Acknowledgments

The authors would like to thank Elizabeth Webster, Dr. Robert Rawle, and Prof. Peter Kasson for helpful discussions. K.N.L. was supported by an NSF predoctoral fellowship. This work was supported by National Institutes of Health Grant R35 GM118044 to S.G.B.

## Supporting Citations

Reference (36) appears in the Supporting Material.

